# THE REACTION TIME COSTS OF TRAJECTORY PLANNING

**DOI:** 10.1101/2022.12.21.521385

**Authors:** Gil Feinstein, Anat Shkedy-Rabani, Lior Shmuelof

## Abstract

Issuing a goal-directed action requires specifying the goal of the action as well as planning the hand trajectory to obtain it. Accumulating results suggest that planning a straight point-to-point trajectory is more efficient and likely to involve simpler optimization process compared to the planning of trajectories with more complex shapes (e.g., curved trajectories). We sought to find evidence for the qualitative difference between the two planning modes through the investigation of reaction times (RT) in a pointing task performed with the wrist. In experiment 1, 18 subjects performed delayed straight and curved via-point reaching movements to arrays of 2 or 8 targets. Both trajectory type and number of possible targets affected RT. In experiment 2 (N=14), we demonstrate a switching cost between the issuing of the two types of trajectories, irrespective of changes in target position. Unexpectedly, trajectory type did not affect RT in experiment 2, likely due to the lack of target pre-cuing in experiment 2. Our results suggest that the planning of curved and straight trajectories differ in their memory load during pre-planning and requires a time-consuming update of the motor commands when switching between straight and curved plans.

## Introduction

The cognitive processes proceeding the issuing a movement are typically separated into two; goal selection (the “what”), and motor planning of the action to obtain it (the “how”)(Wong et al. 2015). These two stages were identified and studied using reaction time (RT) tasks. For example, when studying goal selection, the number of possible targets was shown to affect RT in a logarithmic manner (Fischman 1984; Hick 1952). The RT costs associated with motor planning were shown through an increase in RTs to movements that are composed of multiple sub-goals in reaching (Henry and Rogers 1960) or in speech (Sternberg et al. 1978), and when two actions are performed simultaneously or in close temporal proximity (e.g., Psychological Refractory Period, (Pashler 1994)).

Motor planning costs are not associated only with the compositionality of the action (number of sub-goals), but also with the shape of the trajectory. The measures of complexity of trajectory are not established, partially because the nature of the planning process is still debated (Ben-Shaul et al. 2004; Sakai et al. 2003; Zimnik and Churchland 2021). Nevertheless, shape planning-associated RT costs are expected since it has been shown that the shortening of preparation time affect various properties of curved trajectories (Kohen et al. 2017), and that in an obstacle avoidance task, trajectories with more inflection points evoke longer RTs (Wong et al. 2016).

Another definition of trajectory complexity is based on its deviation from a straight line. Point to point reaching movements are characterized by planning of straight trajectories, even at the expense of a complicated inter-joint coordination patterns (Morasso 1981). This characteristic could be the outcome of solving a cost function aiming to optimize the smoothness of the movement or torque changes (Flash and Hogan 1985), or solving a visual constraint (Flanagan and Rao 1995). The tendency to plan straight reaching trajectories is likely to decrease the planning duration of straight trajectories due to increased planning efficiency following practice (Maslovat et al. 2011) or due to the restricted search space. We hypothesize here that when task’ s constraints, such as via-points or obstacles, will require the planning of curved trajectories, the planning will qualitatively differ from the planning of straight trajectories.

This suggested differentiation between planning of straight and curved trajectories could lead to an explicit differentiation where planning of curved and straight trajectories will be considered different tasks, thereby eliciting switching costs when planning a straight trajectory after a curved one (or the other way around) (Jersild 1927; Rogers and Monsell 1995). We reason that if there is a qualitative difference between the planning strategies of straight and curved trajectories, switching between the planning of straight and curved trajectories will be associated with increases in RTs.

To examine the costs of trajectory planning, we investigated the RT costs associated with the planning of curved compared to straight trajectories and the time costs associated with shifting between them. In Experiment 1, we examined the effect of action selection and motor planning using pre-cueing, under the hypothesis that the number of pre-cued targets will affect RT due to increased demands on action selection, and that pre-cuing of curved trajectories will be associated with increased RT compared to straight trajectories due to increased motor planning demands. In experiment 2, we examine the switching costs between planning and executing straight and curved trajectories in a choice reaction time task. We hypothesized that planning of straight and curved trajectories repetitively will require less RT than planning a curved trajectory after a straight trajectory and vice versa.

## Methods

32 healthy right-handed subjects (aged 22-28, 12 males and 20 females) participated in the experiments that were approved by the local ethics committee, after signing an informed consent form. Subjects were randomly assigned to one of two experiments and were paid approximately 10$ per hour (experiment 1 – N=18, experiment 2 – N=14).

### General approach

All subjects performed an instructed reaching task by moving a computer screen cursor with their right wrist (Fig. 1). The task goal was to guide the cursor from the origin point through a via-point to the chosen target. The via-points were designed to control the type of trajectory that subject was required to perform (via-points between the origin point and the target for straight trajectories and via-points shifted to the left for curved trajectories).

**Figure 1.**
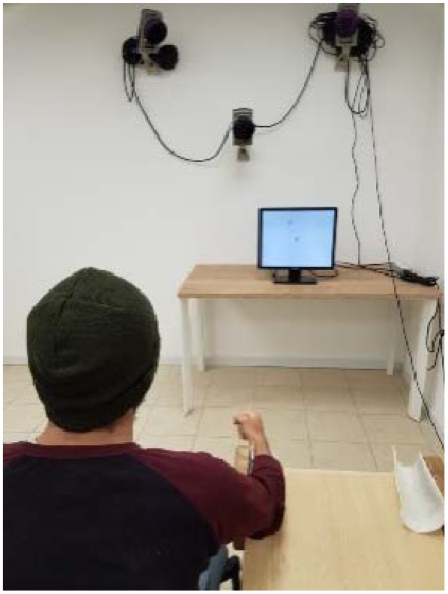
A subject positioned in front of a computer screen. A spherical reflective marker taped on the subject’ s index finger’ s proximal interphalangeal joint (knuckle) is detected by the three Qualisys infrared cameras. The arm of the subject was splinted to a table; his fingers were bent to the arm in a fist. The subject controlled the position of the cursor on the screen by flexing/extending and abducting/adducting his wrist joint.

### Apparatus

Subjects sat 150 cm from a desktop computer screen (19”, DELL 1913S). 3 Qualisys Oqus 500 infrared cameras (Gothenburg, Sweden) recorded the position of a spherical reflective marker taped on the subject’ s index finger’ s proximal interphalangeal joint (knuckle), at a sampling rate of 100 Hz (Fig. 1).

### Task

Subjects moved a cursor between the origin point and the target, through a via-point. The origin point was located at the center of the screen, colored in gray and its radius was 20 pixels. The via-points were indicated by lines (80 pixels length), that subjects had to run over on the way to the targets. The via-point was positioned halfway between the origin and the target. Targets radius was 30 pixels and was located at a distance of 1/4 of the screen size at different angels relative to the origin point (see task for each experiment). In both experiments, the cursor disappeared following movement initiation (at a distance of 10% from the center of the origin point to the target) in order to minimize online corrections and online planning. After every trial, subjects were given two feedback points – one indicating their position at the radius of the via-point and the other at the radius of the target. The target and the via-point were colored according to the performance of the subjects (green indicating a hit and red a miss).

### Experiment 1

Experiment 1 included 4 experimental conditions (two-by-two experimental design), presented in blocks. The two factors were the type of the trajectory (straight or curved, determined by the position of the via-point) and the number of possible trajectories (2 or 8) (Fig. 2A). Before the beginning of the experiment, subjects performed a training block of 50 trials. Subsequently, subjects performed 8 blocks (2 per condition, 50 trials in each block). Each trial was composed of 3 phases (Fig. 2B): 1. Optional targets presentation (pre-cueing, 1.5 sec)-optional targets and their associated via-points appeared on the screen (two straight/two curved/eight straight/eight curved) 2. Go signal (cueing, 1.5 sec) - all targets and via-points disappeared except one target and its associated via-point that were colored in blue, indicating the subject to move to that target. 3. Feedback (as described above, 1 sec). In this experiment, 2 different gray intensities were used for coloring targets and via-points that are close to each other to emphasize the association between the via-point and targets. Targets were separated by 45° (Target 1 was positioned at 22.5°). For each target there were 2 optional via-points: in the straight condition, the via-point was presented exactly on the midline connecting the target and the origin, and in the curved condition - 30° to the left of the midline. Subjects had to move to the indicated target through the presented via-point. In all trials, the target and via point were specified to the subjects at the Go signal.

**Figure 2:**
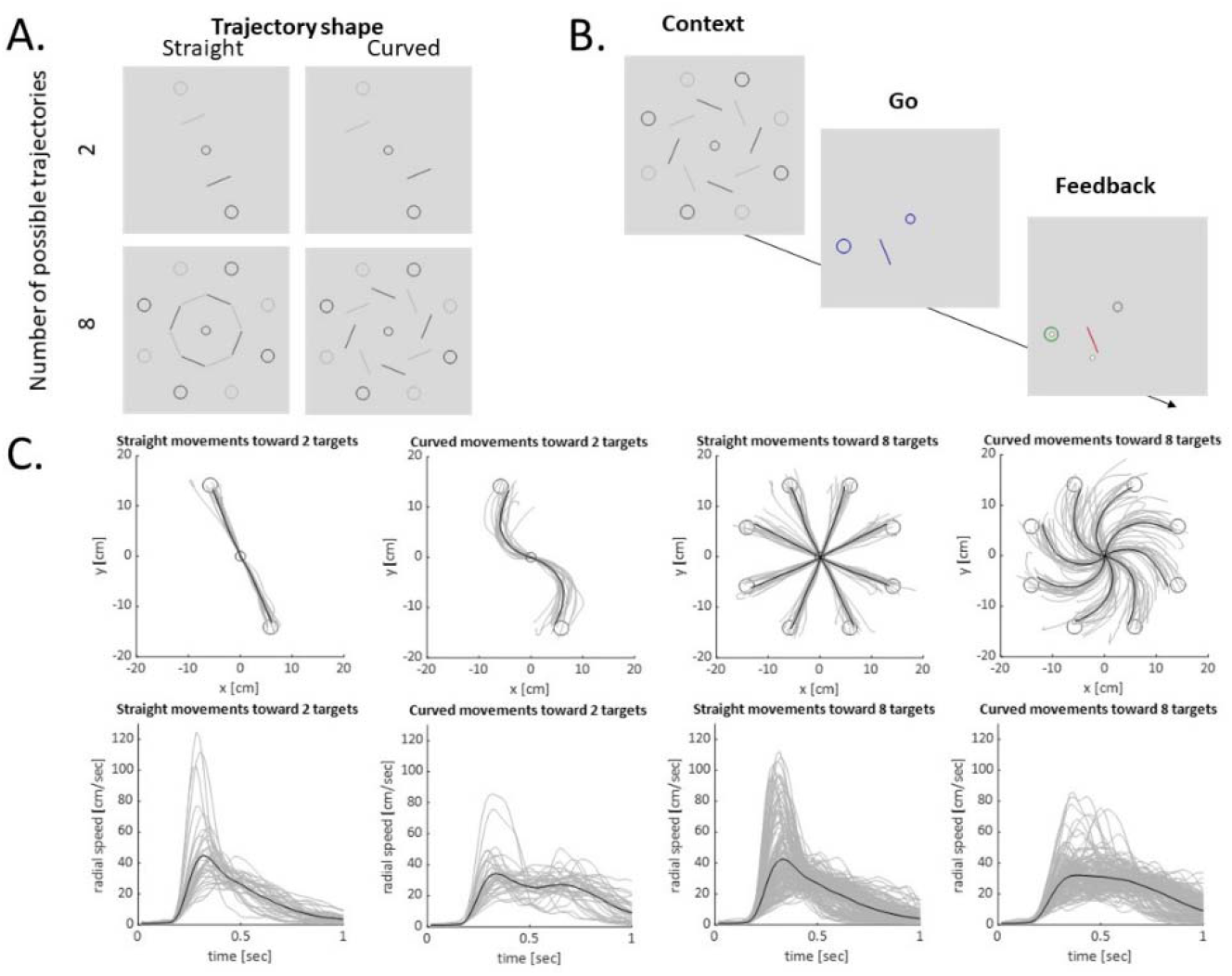
Experiment 1. A. Description of the 4 possible conditions (2 straight optional trajectories/2 curved optional trajectories/8 straight optional trajectories/8 curved optional trajectories). B. Description of the 3 phases in a trial. C. Averaged trajectories from the four conditions (top) and their velocity profiles (bottom). Gray lines represent single subject average trajectories to each target. Black line represent the grand average.

### Experiment 2 – switching cost

The purpose of this experiment was to study the switching costs between planning and execution of straight and curved trajectories. Experiment 2 included 3 experimental conditions, presented in blocks (Fig. 3A): 1. Fixed straight –straight trajectories to 4 possible targets. 2.

**Figure 3:**
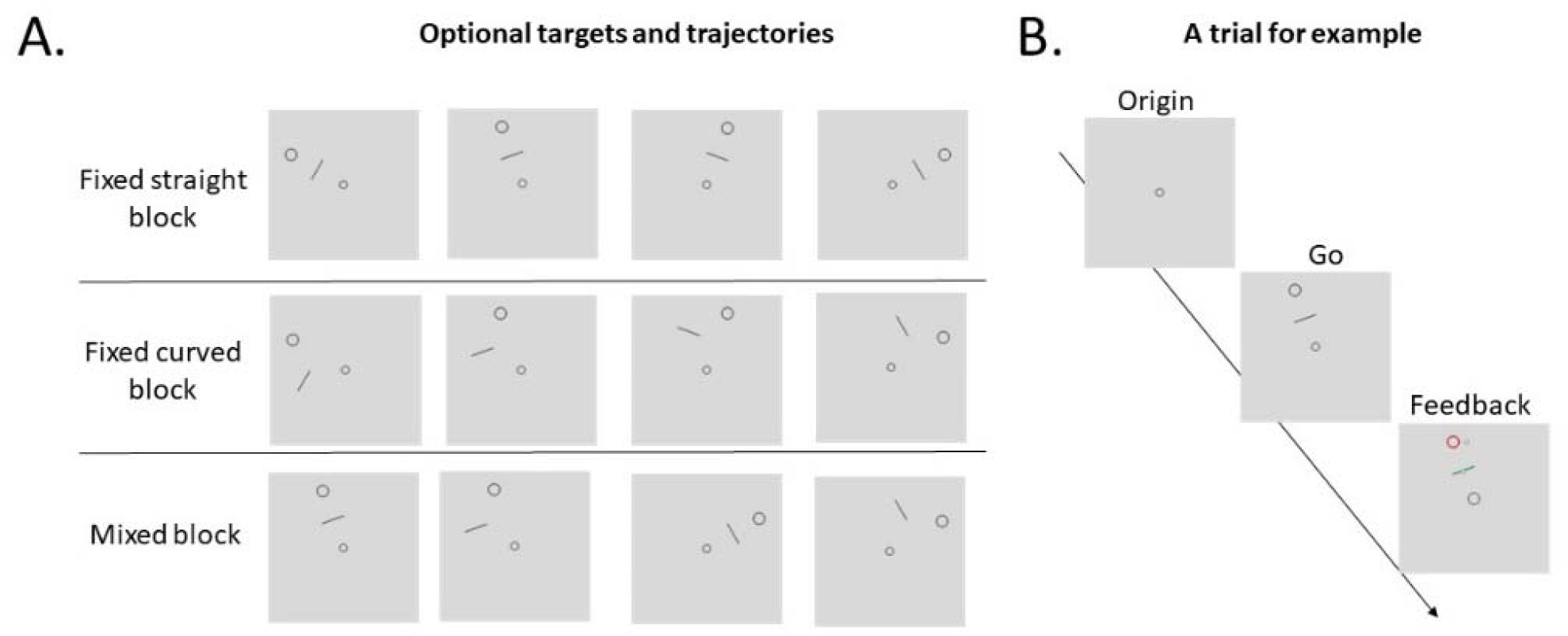
A. All possible targets and via-points in each block. B. The presentation during the 3 phases of a trial: Return to the origin, Appearance of the target (the Go signal) and Feedback presentation.

**Figure 4:**
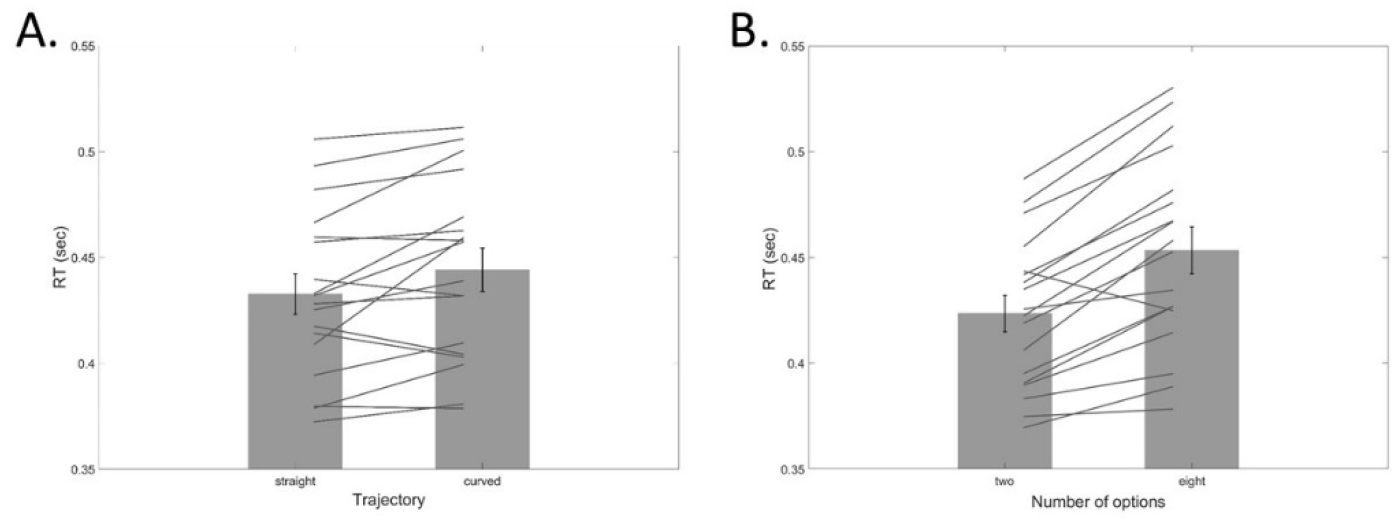
The effect of type of trajectory (A) and number of optional trajectories on RT (B). Gray lines represent performance of individual subjects. Error bars represent SEM.

Fixed curved –curved trajectories to 4 possible targets. 3. Mixed – Both straight and curved trajectories to 2 possible target (4 possible trajectories). Before the beginning of the experiment, subjects performed a training block with 20 trials. Subsequently, subjects performed 8 50-trials blocks. Two sequence variants were used across subjects: (1) fixed straight, fixed curved, 4 mixed, fixed curved and fixed straight and (2) fixed curved, fixed straight, 4 mixed, fixed straight and fixed curved. Each trial was composed of 3 phases (Fig. 3B): 1. Origin (1.5 sec) – after the completion of the previous trial, subjects received feedback of cursor’ s position and an instruction to bring the cursor to the origin point and wait. 2. Go signal (cueing, 1.5 sec) - the chosen target (and its via-point) appeared, and subjects were required to move the cursor from the origin to the target through the via-point. 3. Feedback (as described above, 1 sec). There were 4 optional target positions in this experiment (30°, 70°, 110°, 150°). Each target could be presented with one of two optional via-points (on the midline connecting the origin and the target for straight trajectories and shifted 45° to the left for curved trajectories). The general structure of the experiment was designed based on previous switching cost experiments (Meiran 2014).

Importantly, the target for the reaching was presented only at the Go signal. In the mixed condition, the 30° and 110° targets were presented.

### Power analysis

Sample size was computed based on the results of Wong et al., (Wong et al. 2016), as presented in experiment 1, where an estimate of 16 ms difference between the planning of a straight and curved trajectories is reported. Variability estimate was taken from the same experiment from the T^plan^ condition (SD = 22.31ms), indicating an effect size of 0.73. To detect such an effect with a power of 0.8, data from an estimate of 14 subjects is required.

### Data analysis

RT was defined as the time it took subjects to reach 5% of the movements’ first velocity peak. This approach was chosen to account for differences in the velocities between the two types of trajectories. After calculating RT for each trajectory, the mean RT and the standard deviation were calculated. Trials with anomalous RTs were removed (3 SDs). Statistical analyses were made using SPSS and JASP. Repeated measures ANOVA and paired t-tests were used for statistical inference. Accuracy was a binary measure which was based on the overlap between the cursor and the via point and the target.

## Results

### Experiment 1

To examine the effect of trajectory shape and number of possible trajectories on motor planning duration, we compared the average RTs in each condition using a repeated measures ANOVA analysis.

| <u>Block</u> | <u>Mean RT</u> | <u>SD</u> |
| --- | --- | --- |
| 2 possible straight targets | 0.416 | 0.035 |
| 2 possible curved targets | 0.431 | 0.039 |
| 8 possible straight targets | 0.449 | 0.046 |
| 8 possible curved targets | 0.458 | 0.048 |

Straight trajectories were issued 15 ms faster than curved trajectories (F(17,1) = 8, p = 0.012, partial η^2^ = 0.32) and trajectories that were selected out of 2 options were issued 30 ms faster than trajectories that were selected out of 8 options (F(17,1) = 44.4, p < 0.001, partial η^2^ = 0.723). There was no interaction between the effect of number of possible trajectories and the effect of trajectory type (F(17,1) = 1.27, p = 0.275) (Table 1). The shorted RT for the straight trajectories indicates that either their motor planning was shorter than the planning of the curved trajectories, or that their motor planning began during the pre-cueing phase, whereas the plan of the curved trajectories was not processed to the same degree during the pre-cueing phase.

Curved trajectories were less accurate than the straight trajectories. The proportion of trajectories where both the target and the via-point were reached was higher in the straight compared to the curved conditions (F(1,13) = 5.39, p=0.037).

### Experiment 2

To further examine the difference between planning of straight and curved trajectories, we examined the switching costs between the planning of straight and curved trajectories.

Subjects were asked to plan and execute straight and curved reaching trajectories, either in a fixed block where only one type of trajectory was required or in a mixed block where both types were required. In both cases, the target of the trial appeared with the go cue, preventing pre-planning of the trajectories. Trials were then divided into 8 categories according to their shape (straight/curved) and the repetition context of the trajectories (fixed, mixed after a switch, mixed after a one no-switch trial, and mixed after two no-switch trials).

| <u>Sub-group</u> | <u>Description</u> | <u>Mean RT</u> | <u>SD</u> |
| --- | --- | --- | --- |
| <u>Straight trajectories</u> |  |  |  |
| S | fixed block | 0.404 | 0.033 |
| S1 | mixed block following a curved trajectory | 0.434 | 0.042 |
| S2 | mixed block following S1 trajectories | 0.420 | 0.040 |
| S3 | mixed block following S2 trajectories | 0.415 | 0.039 |
| S – all | All straight trajectories together | 0.418 | 0.010 |
| <u>Curved trajectories</u> |  |  |  |
| C | fixed block | 0.405 | 0.044 |
| C1 | mixed block following a straight trajectory | 0.428 | 0.043 |
| C2 | mixed block following C1 trajectories | 0.423 | 0.043 |
| C3 | mixed block following C2 trajectories | 0.420 | 0.046 |
| C – all | All curved trajectories together | 0.414 | 0.009 |

Repetition context had a significant effect on RT (Fig. 5A) (F(3,78) = 38.372, p<0.001, partial η^2^ = 0.596). A movement that was issued in a fixed block (straight or curved) required the shortest RT (404 and 405 ms, respectively), a movement that was issued in a mixed block after a switch (straight after curved or curved after straight) required the longest RT (434 and 428 ms respectively). RT decreased as subjects repeated trajectories of the same shape within the mixed block (F(2,52) = 31.56, p < 0.001, partial η^2^ = 0.548). The third repetition of a trajectory within the mixed block required longer RT than repeating that trajectory in the fixed block (t(26)= 3.057, p=0.005 for the straight trajectories and t(26)= 2.452, p=0.021 for the curved trajectories).

**Figure 5:**
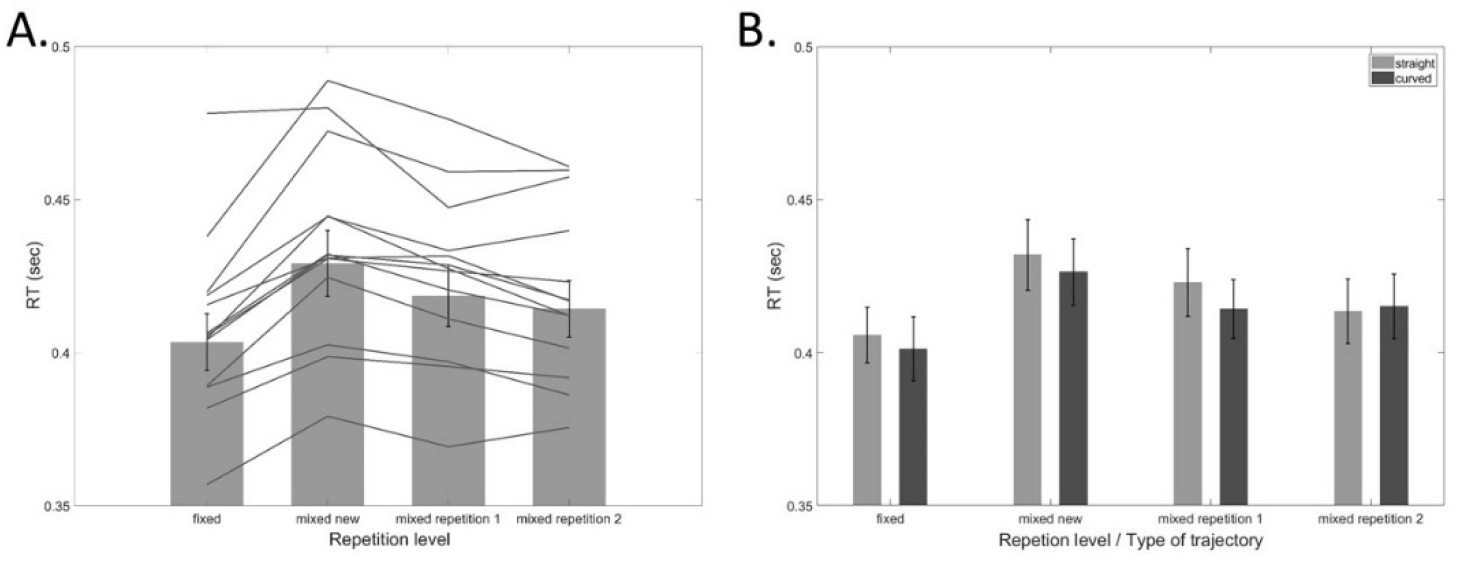
A. Averaged reaction times at 4 repetition levels, highlighting the difference between the fixed condition, where subjects performed only straight or curved trajectories, and mixed conditions where they made both straight and curved trajectories in the same blocks. Gray lines represent performance of individual subjects. B. Averaged reaction times for repetition levels separated for straight (light gray) and curved (dark gray) trajectories. Error bars represent SEM.

Interestingly, evidence of the motor planning effect with respect to the type of trajectory was not found (RTs of straight and curved trajectories did not differ, F(1,26) = 0.038, p=0.847) (Fig. 5B). This result suggests that the observed motor planning effect in experiment 1 is driven by differences in the pre-cuing processing of both trajectories. Under pre-cueing conditions, subjects planed straight trajectories to a greater extent than curved trajectories. This result also indicates that asymmetry in switching costs cannot be investigated in this experiment.

Like experiment 1, accuracy was decreased for the curved trajectories. Subjects reached the via-point and the target significantly more during the straight trials compared to the curved trials (F(1,13)= 14.86, p<0.001). The context of the trial (fixed/mixed) did not affect accuracy (p>0.2).

## Discussion

We report two distinct effects of motor planning on reaction times. First, we show that planning curved trajectories led to increased RT compared to straight trajectories (irrespective of the number of possible trajectories), and second, we report a switching cost between straight and curved of trajectories, which provide further support for the existence of a time-consuming trajectory planning process.

Trajectory type did not affect RT in experiment 2. We suggest that this apparent discrepancy between experiments 1 and 2 stems from the pre-cueing of the targets in experiment 1. In the presence of explicit knowledge about the upcoming targets (such as in pre-cueing designs), planning of movement’ s trajectories begins before the go cue, presumably to improve performance or decrease RT (simple reaction time tasks). In the absence of explicit knowledge prior to the go signal, trajectory planning starts only after the ‘go signal’ (choice reaction time tasks). Our results indicate that trajectory type effect happens only in pre-cueing conditions, suggesting that RT costs reflect a strategic reaction to pre-planning conditions, rather than a direct outcome of the time difference between the generation of a straight and a curved motor plans. The switching cost, on the other hand, was observed without explicit knowledge of the repeatability of the trajectory. Thus, it could reflect a cost of an execution process rather than a planning process. The fact that switching was seen in conditions where deliberate pre-planning is unlikely to occur suggests that it is an outcome of mechanism that enhanced repetition based on a memory of the previous command (caching) (Jax and Rosenbaum 2007).

A recent perspective highlights the fact that certain aspects of the movement are not planned before the GO cue, even when targets are pre-cued, presumably due to the high cognitive load that is associated with maintaining these plans. Thus, even when the target is pre-cued (like in experiment 1), the complexity of the task still affects its RT(Klapp and Maslovat 2020). This result is consistent with the results of experiment 1 that showed that even though the targets were shown before the GO signal, and could be potentially planned before the go cue, the number of options and their shape affected the RT. Considering the previous results from the simple reaction time task, it suggests that both action selection and shape planning cannot be fully conducted before the GO cue. In experiment 2, that was conducted under a choice reaction time design, when straight and curved trajectories appeared repeatedly (fix block), they required comparable RTs. Taking the trajectory shape RT results of the two experiments together, our results suggest that while straight trajectories are amenable to pre-planning, curved trajectories are less so. When preplanning is impossible – such as in the case of choice RT task, these two shapes require the same RT, but when preplanning is allowed (experiment 1), preplanning of straight trajectories is greater than preplanning of curved ones.

Interestingly, the RTs in the fixed block in experiment 2 were shorter than the RTs in the 2 options condition in experiment 1 (404 ms vs 416 ms for the straight conditions and 405 ms vs 431 ms for the curved condition). The interpretation of this effect is challenging since the two experiments differed both in the pre-cueing and in the action selection demands. It could be that pre-cuing of multiple targets in Experiment 1 led to some degree of planning of multiple trajectories, so that subjects had to apply a time-demanding suppression of the un-cued plans following the go cue. Alternatively, executed motor plans in experiment 2 could be saved (cached) across trials, thereby leading to (1) reduced RT when compared to a condition where subjects pre-planned the trajectories on a trial-by-trial basis (experiment 1) and (2) a switching cost when the required trajectory changed (experiment 2). Thus, deliberate pre-planning (experiment 1) may not be an effective strategy when compared to a ‘ minimum planning’ mode, in the context of repeated stimuli.

The ability to plan a complex trajectory following a demonstration or an abstract instruction is impaired in patients that suffer from ideomotor apraxia, which is associated with damage to the left parietal and premotor areas (Buxbaum et al. 2014). In a recent study, Wong et al. have shown that apraxia patients can imitate straight point to point trajectories but show deficits when asked to imitate or copy complicated shapes (such as a sine wave) (Wong et al. 2019). These neuropsychological results support our claim regarding the dissociation between planning of straight and curved trajectories and suggest that the impaired ability in ideomotor apraxia patients may be the time-consuming trajectory planning function that is described in our experiments. It also raises the conjecture that imitation resembles the process of planning a trajectory based on spatial constraints (such as based on via-points and obstacles), suggesting that both processes involve generating a motor plan based on an abstract representation of a trajectory.

Reaction time differences are used here as an indirect estimate of the mental processing that takes place before the initiation of the movement (Posner 1978). Importantly, motor planning does not stop at movement initiation and continues during its execution (Chen-Harris et al. 2008; Todorov and Jordan 2002). In fact, there is a tradeoff between RT and online planning/movement quality (Kohen et al. 2017; Song and Nakayama 2008). This trade-off may also affect the performance of subjects in our experiments; the curved trajectories were less accurate than the straight ones, suggesting that the quality of their planning was decreased in order to reduce the RT. This interpretation of the results indicates that the RT costs of trajectory planning are an underestimate of the planning time that is required to plan a trajectory with a complex shape.

We hypothesized that the effect of number of optional targets is linked to the selection of motor goal whereas trajectory planning happens later in time. This hypothesis was supported by the lack of an interaction effect in experiment 1. Nevertheless, the average RT difference between straight and curved trajectories in the condition of 2 possible targets (15 ms) was longer than the same difference in the condition of 8 possible targets (8 ms). We cautiously suggest that the number of possible targets can indirectly affect the trajectory planning process, consistent with the idea that the process of issuing a goal-directed action is optimized as a whole (Wong and Haith 2017). When the amount of options increase beyond a certain limit, the motor system may stop pre-planning each possible trajectory serially, and trigger a strategy of postponing pre-planning overall. In our case, it could be that the pre-planning strategy was abandoned in some of the 8-target trials, thereby leading to reduced trajectory shape effect. Future experiments should examine, using RT and kinematic analyses, whether the reported difference between the planning of straight and curved trajectories reflects a secondary, pre-planning strategy effect, rather than a direct effect of trajectory planning effort on reaction time.

## Acknowledgments

We thank Or Dezachyo and Uri Monsonego for their assistance in collecting the data. We thank John W. Krakauer and Nachshon Meiran for valuable comments on the manuscript. This research was supported by the Helmsley Charitable Trust through the Agricultural, Biological and Cognitive (ABC) Robotics Initiative and by the Marcus Endowment Fund both at Ben-Gurion University of the Negev, by the United States-Israel Binational Science foundation grant 2015327, and by ISF grant 607/16 for Lior Shmuelof.

